# Computational design of genes encoding completely overlapping protein domains: Influence of genetic code and taxonomic rank

**DOI:** 10.1101/2020.09.25.312959

**Authors:** Stefan Wichmann, Siegfried Scherer, Zachary Ardern

## Abstract

Overlapping genes (OLGs) with long protein-coding overlapping sequences are often excluded by genome annotation programs, with the exception of virus genomes. A recent study used a novel algorithm to construct OLGs from arbitrary protein domain pairs and concluded that virus genes are best suited for creating OLGs, a result which fitted with common assumptions. However, improving sequence evaluation using Hidden Markov Models shows that the previous result is an artifact originating from dataset-database biases. When parameters for OLG design and evaluation are optimized we find that 94.5% of the constructed OLG pairs score at least as highly as naturally occurring sequences, while 9.6% of the artificial OLGs cannot be distinguished from typical sequences in their protein family. Constructed OLG sequences are also indistinguishable from natural sequences in terms of amino acid identity and secondary structure, while the minimum nucleotide change required for overprinting an overlapping sequence can be as low as 1.8% of the sequence. Separate analysis of datasets containing only sequences from either archaea, bacteria, eukaryotes or viruses showed that, surprisingly, virus genes are much less suitable for designing OLGs than bacterial or eukaryotic genes. An important factor influencing OLG design is the structure of the standard genetic code. Success rates in different reading frames strongly correlate with their code-determined respective amino acid constraints. There is a tendency indicating that the structure of the standard genetic code could be optimized in its ability to create OLGs while conserving mutational robustness. The findings reported here add to the growing evidence that OLGs should no longer be excluded in prokaryotic genome annotations. Determining the factors facilitating the computational design of artificial overlapping genes may improve our understanding of the origin of these remarkable genetic constructs and may also open up exciting possibilities for synthetic biology.

## Introduction

The triplet nature of the standard genetic code and double-stranded configuration of DNA together enable more than one protein to be encoded within the same nucleotide sequence in different reading frames. This property of the code has long been known to be utilised in viruses [1,2] and there is increasing evidence for overlapping encoding in other organisms [3, 4, 5], including many genes fully embedded within other coding sequences in alternate reading frames [6]. While a mutation in a stop codon can easily create a short, trivial overlap in neighbouring genes as a chance event, longer, non-trivial overlaps should only be maintained in a genome if the overlapping region encodes a part of the protein essential for its function for both genes. There are a few hypothetical reasons why genes might overlap, and the evidence for functional antisense overlaps in prokaryotes has been discussed in a recent review [7]. While the reduction of genome size is particularly relevant only for some viruses [8, 9], it has also been studied in bacteria [10]. Effects on gene regulation [11] conceivably could affect all organisms, for instance there is the possibility of co-expression of same-strand overlapping genes (OLGs) with the mother gene, given that they are potentially expressed from the same mRNA. Genes within an antisense overlapping pair could also influence each other, for instance in a way similar to what has recently been termed a “noncontiguous operon”, where genes in antisense to each other are nonetheless co-expressed as an operon [12]. Other proposed benefits of overlapping genes relate to templating structure based on the existing ‘mother gene’, namely, for genes directly in antisense (“-1 frame”), the creation of proteins with a complementary polarity structure to the gene on the antisense strand [13, 14, 15] or, in the case of sense overlaps, a similar hydrophobicity profile [16]. Overlapping open reading frames may play an important role in the origin of *de novo* genes, exploring new territory in the total space of sequences and functions [17, 18, 19, 20]. While most currently extant OLGs are not taxonomically conserved and therefore appear to be evolutionarily young [51], one claimed example of an ancient OLG pair is comprised of the two classes of aminoacyl-tRNA synthetases which can be encoded in an overlapping manner [21, 22, 23].

Despite the many possible effects of overlapping genes (OLGs), they are generally not considered a significant phenomenon outside of viruses, due perhaps to perceived difficulties in their evolution for some or all reading frames [24, 26]. The idea that they have been more widespread has long been theorized [25, 26, 56]. As a consequence, most gene prediction algorithms still exclude non-trivially overlapping genes [27], especially outside of bacteriophages and other viruses. The NCBI rules for annotation of prokaryotic genes do not allow genes completely embedded in another gene in a different frame without individual justification [28]. Even in viruses, relatively few overlapping genes have been annotated, particularly antisense gene pairs, although more are regularly being discovered including, for instance, in the pandemic viruses HIV and SARS-CoV-2 [2,29,50].

A recent study [30] quantified the difficulty of constructing OLGs by picking random pairs of protein domains and rewriting them so as to overlap, with an algorithm minimizing the amino acid changes in each domain. This is a new approach, as previous studies tried to create overlaps without changing the amino acid sequence of the two genes, which resulted in either a very limited overlap length [31] or could only be done for very specific genes [32]. They found that, remarkably, 16% of 125250 arbitrary protein domain pairs were able to successfully overlap in at least one of the 3 reading frames they investigated, and one of two positions tested. Virus domains were much more likely to create putatively functional overlaps than domains from prokaryotes or eukaryotes, as determined by BLAST searches of the SWISS-PROT database. This result suggests that creating overlaps is not as difficult as might be expected, implying that an abnormally high threshold of evidence as compared to other gene types should not be required for verifying their existence. This high success rate also opens up many possibilities for synthetic biology. For instance, mutations in overlapping regions are expected to be more deleterious on average, so an artificial genome with many OLGs is not only smaller but also expected to be more stable over time on a population level, as mutations are more likely to be strongly selected against. A recent method for stabilizing synthetic genes [33], where an arbitrary ORF was constructed to overlap a gene of interest and was concatenated with an essential gene downstream, could be taken a large step forward by overlapping whole genes thereby creating a system where not only ‘polar’ mutations are selected against but also more minor mutations, if they also affect the mother gene. Genome size has become a significant limiting factor for biomolecular computing, in which genetic programs are inserted into cells [34]. Existing compression methods [35] could be greatly improved by using OLGs, making more complex systems possible. In this context a well designed stable synthetic genome could include fail-safe measures, such that faulty genetic programs would shut down.

Here the algorithm provided in [30] is used but improved in the evaluation of the constructed sequences as the analysis in the previous study has some weaknesses resulting in incorrect claims. Determining whether an artificial sequence has a specific function from its amino acid sequence only is a very hard problem and not possible today. Progress is being made in predicting the protein structure from amino acid sequence [53], but protein structure does not determine function as essential binding sites can be rendered useless if the amino acid is changed while not changing the overall protein structure. Ultimately only experiment can definitively determine the function of a given amino acid sequence. In order to aid the design of expensive experimental setups however, it can at least be determined bioinformatically how similar an artificial sequence is to sequences with known functions. In this study the artificially designed sequences are compared to their original sequences in terms of amino acid identity, amino acid similarity, Hidden Markov Model profile and secondary structure in order to determine the impact of OLG construction and which sequences are potentially functional.

Firstly, the details on how some technical artifacts arose are explained and how to avoid them. In order to further improve the analysis Hidden Markov Models rather than BLAST is used in this study. While the previous study [30] tried to estimate an upper limit of how many domains can be successfully overlapped in at least one reading frame and position, here the average success rate for OLG construction is determined instead, which is more relevant in relation to both understanding constraints on the formation rate of naturally occuring OLGs and in assessing the likelihood of successful synthetic creation of OLGs. These results in one sense give an upper estimate of the ease of creating overlaps as the difficulty of obtaining an overlapping gene pair naturally is not directly addressed here. On the other hand, overlapping functional domains directly is a “worst case scenario” as there is some evidence that the critical functional domains of one protein in an OLG pair tend to overlap less constrained regions of the other protein [36], and this segregation is also intuitively plausible. In order to estimate the difficulty of achieving overprinting naturally, the minimal number of nucleotide changes needed to create the OLG sequence is determined. Whether functional domains do in fact overlap in nature, however, deserves further attention.

By expanding the analysis of the previous study [30] from the reading frames ‘+2’,’-1’ and ‘-3’ to all reading frames (see Fig. 1 for reading frame definitions), the observed differences between reading frames can be related to the structure of the standard genetic code. Through constructing OLGs using randomly generated genetic codes it can be studied whether the standard code shows evidence of optimisation regarding OLGs. Using the improved evaluation of the designed OLGs it can be shown that virus genes, surprisingly, are less suited than bacterial and eukaryotic genes to design OLGs.

**Figure 1:**
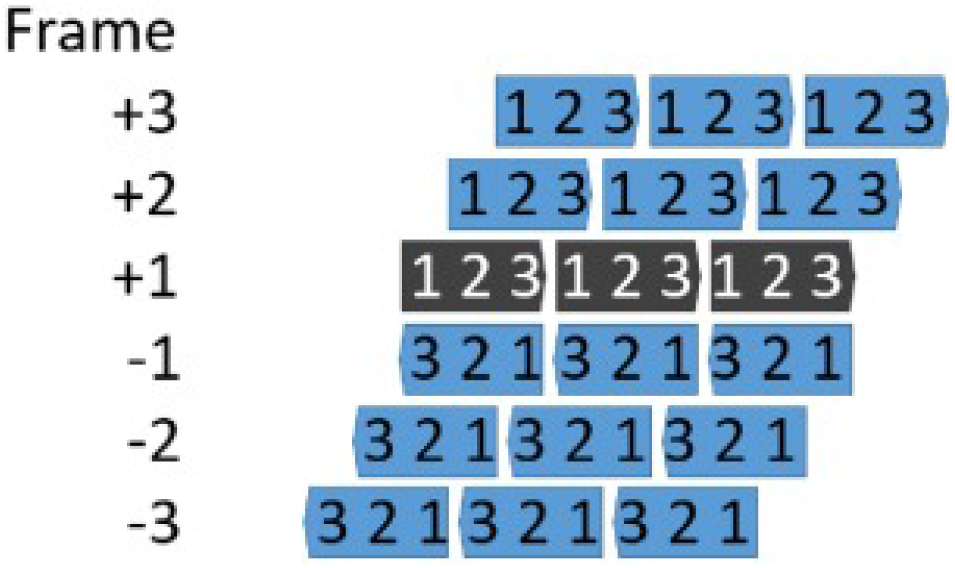
Illustration of the alternative reading frames. The ‘+1’ frame is the standard or reference reading frame and ‘+2’/’+3’ the sense overlaps, while frames ‘-1’ to ‘-3’ are on the anti-sense strand.

## Materials and Methods

### Dataset-Database biases

In [30] constructed sequences were evaluated with a BLAST search against the SWISS-PROT database. If both overlapping sequences had a match to the best hit with at most an e-value of 10^(−10) and a match length of 85%, the overlap was considered successful. However, the initial sequences were picked from the Pfam seed database and it can be shown that most of the chosen sequences are not well represented in the SWISS-PROT database (see left panel in Fig. 2), with the exception of virus genes. In a search against the SWISS-PROT database, identities of over 80% were only found for 15% of the non virus genes, while 70% of the virus genes could be found in this category. A curated set in which all sequences have a 100% match in the SWISS-PROT database but otherwise the same properties has a remarkable 95% success rate for overlaps and the virus vs non-virus difference vanishes (see right panel in Fig. 2). The advantage reported for virus genes is thus fully explained by dataset-database biases. In any case, the extremely high overall success rate obtained should be investigated. Either creating overlaps is indeed unexpectedly easy or the evaluation of functionality used in [30] is not conservative enough. It can be shown that both factors appear to contribute to the surprising result.

**Figure 2:**
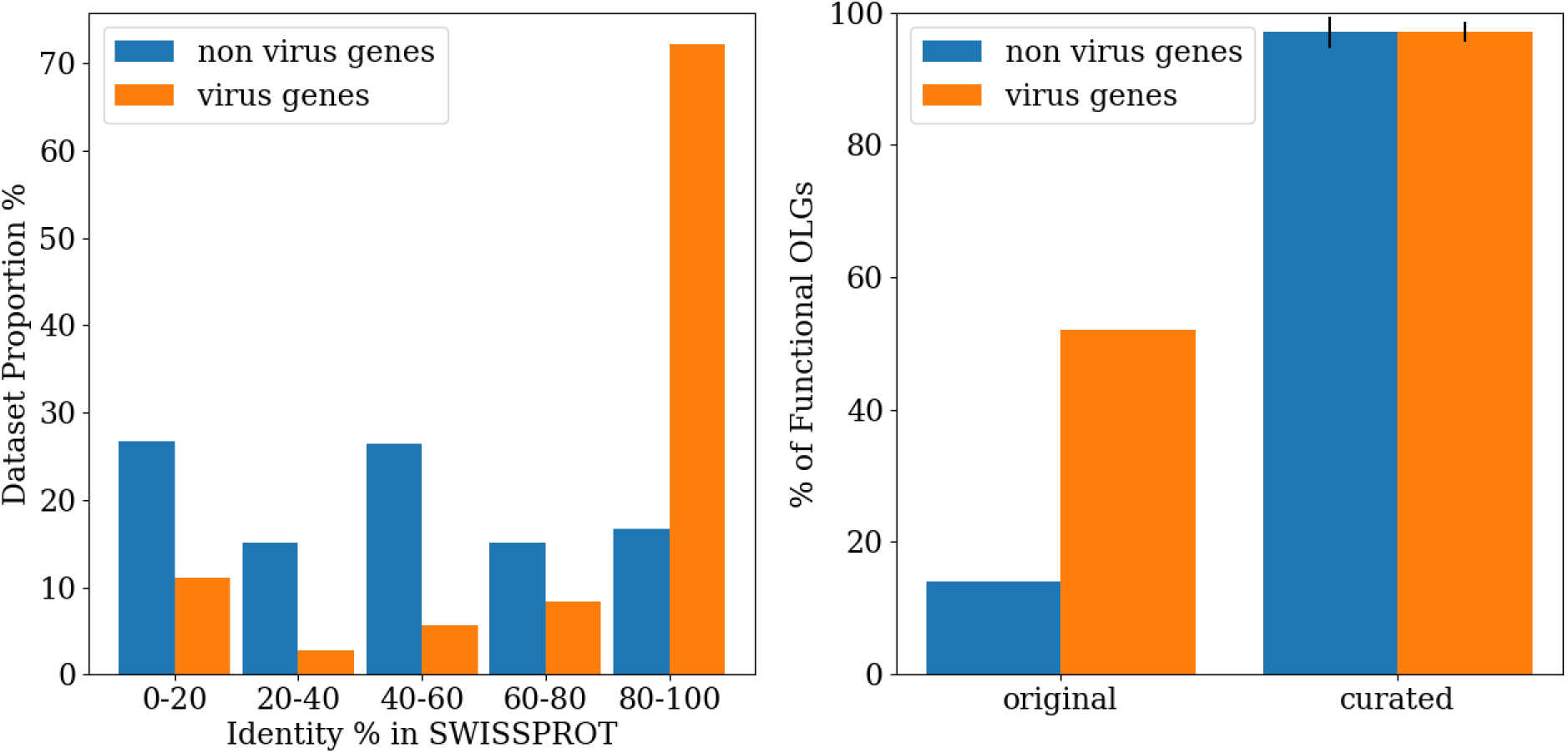
*Left:* Proportions of the dataset used in [30] with different match identities in SWISS-PROT - virus genes from this dataset have a higher average identity to a SWISS-PROT entry than non-virus genes. *Right:* Percentage of functional OLGs for the original dataset used in [30] and the average of 10 curated datasets grouped into virus and non virus genes. In curated datasets all original sequences have an exact match in SWISS-PROT. Each curated dataset has 100 sequences with 70-100 amino acids. The virus versus non-virus difference observed in the dataset of [30] vanishes for the curated datasets.

### Length dependence due to a fixed cutoff

When introducing the minimal number of changes required for two random sequences to fully overlap each other, a similar percentage of each sequence is expected to change. In such a case the e-values of the constructed sequences would be strongly length dependent, as a longer sequence with the same similarity has a lower probability of being found by chance in a database of a given size. When picking datasets with different sequence lengths such a length-dependence can be found in the BLAST evaluation (supplementary Fig. S1). A fixed e-value cutoff cannot adequately evaluate sequences in such a situation as the cutoff value fully determines the result and is chosen arbitrarily. The sequences used in [30] have a length of 70-100 amino acids, and the high success rate for the curated set can be explained by a combination of the sequence length and the choice of the cutoff value.

### Hidden Markov Models combined with the Pfam database solve the cutoff problem

In order to find a reasonable alternative to the fixed e-value cutoff, Hidden Markov Models (HMMs) can be used to score the constructed sequences. Here HMMER3 (v3.2.1) [37] is used to create profiles for each protein domain family in the Pfam database [38] in order to score the constructed sequences. The Pfam database consists of a ‘seed’ database, containing trusted sequences for each family which are used to create HMM profiles, and a ‘full’ database, containing all the sequences of the Uni-Prot database sorted into the different families according to the previously constructed profiles. Here the HMM profiles are also constructed from the ‘seed’ sequences and in order to find the sequence most closely representing the profile all full sequences are tested against the profiles. The highest scoring sequences are used to construct OLGs. The rest of the ‘full’ sequences are used as a comparison for the overlapping region of the constructed sequences. A constructed sequence is judged successful if it has a higher score than a sequence at a defined threshold percentile of the ‘full’ sequences, thereby creating a threshold value which is individual for each protein family. Here results for different threshold percentiles are discussed, while highlighting two particular percentile values. Firstly, the 50th percentile (median), which marks the score of a typical sequence in the protein family. In this analysis, sequences meeting this threshold can not be distinguished from the naturally occurring protein domains and they will be categorised as typical proteins. Since all sequences in the ‘full’ group are naturally occurring sequences, scoring at least as highly as any of these sequences renders a sequence biologically relevant. In order to avoid extreme outliers which may be misclassified, the 5th percentile is used as the biologically relevant threshold.

A relative threshold could alternatively be established with e.g. BLAST by first picking a single sequence as a starting point for construction and also for comparing the rest of the protein family to in order to find the threshold score as described above. In this case however, it is not clear which sequence to choose as a starting point. A randomly picked sequence could be an outlier of the protein family, resulting in unreliable comparison scores and a higher chance of losing function after constructing OLGs. HMMs on the other hand provide a profile reflecting the ‘average’ sequence, which is a better representative for the whole protein family.

Choosing a family-specific threshold value takes care of most of the length dependencies, but in order to be sure and to be able to compare sequences of different lengths, each score resulting from a comparison between a sequence and a HMM profile is divided by the sequence length. Here scores are used instead of e-values, as the latter also depend on the database size, an arbitrary factor in this analysis. Aligning the best sequence with the ‘seed’ sequences using MAFFT (v7.419) [39], weights used for sequence construction can be determined just as in [30]. A more detailed description of the calculation of the weights and their influence can be found below. When studying the influence of a protein family’s taxonomic classification on the construction of OLGs, the ‘seed’ and the ‘full’ database are first filtered by the four major taxonomic groups - archaea, bacteria, eukaryotes and viruses - before creating the profiles and the thresholds. MUSCLE (v3.8.31) [40] was used for realigning the ‘seed’ sequences after taxonomic filtering.

For subsequent analyses, random sets from the ~17000 Pfam families were chosen, with the condition that each family must have at least 10 ‘seed’ sequences and 4 ‘full’ sequences in order for the weights and the thresholds to be reasonably defined. Each dataset consists of 150 families since the variance of the resulting OLG success rate barely declines for larger sets (see supplementary Fig. S2). Fig. 3 summarizes the workflow.

**Figure 3:**
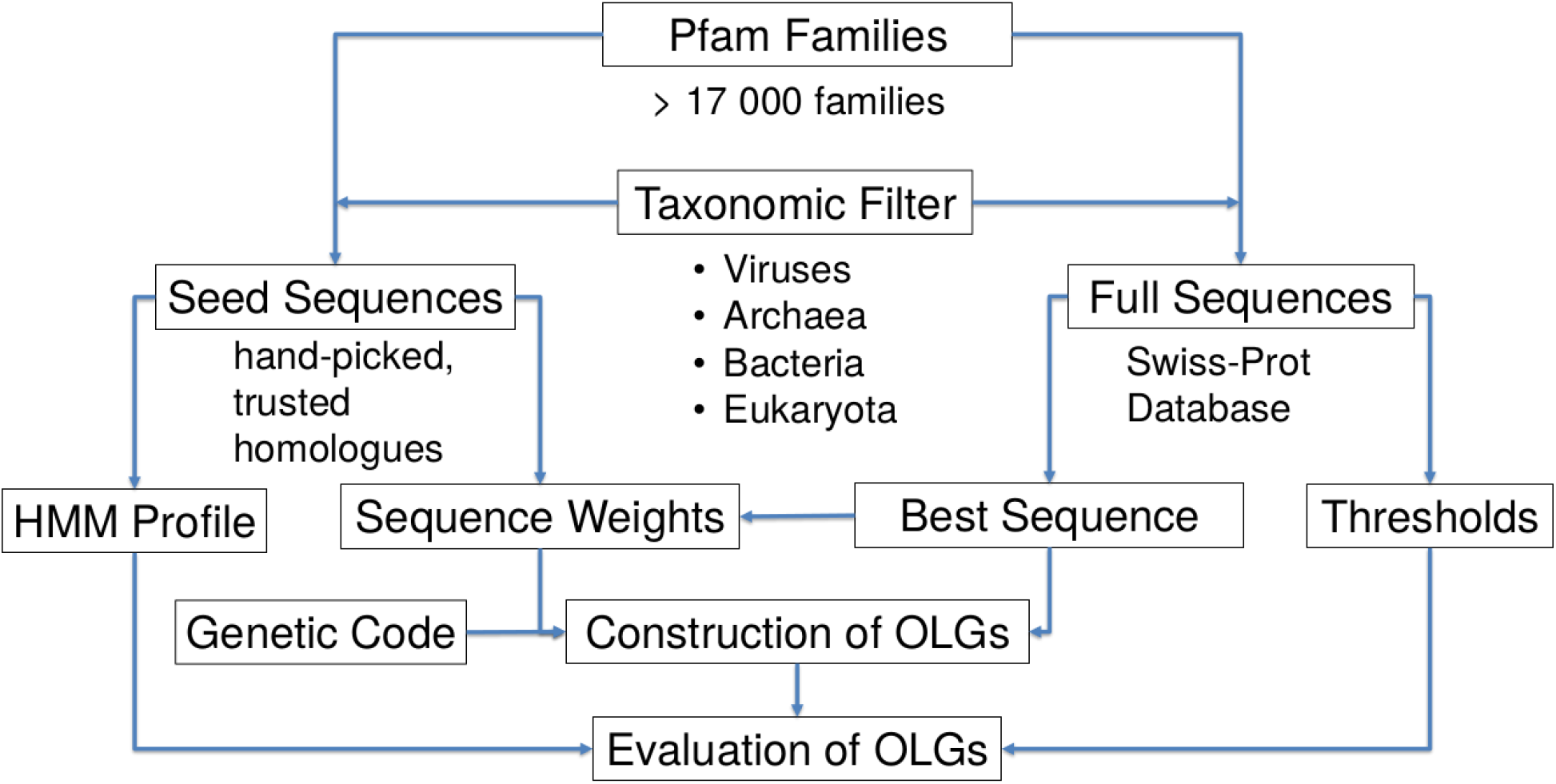
Workflow for OLG construction and evaluation using HMMs and the Pfam database. HMM profiles are constructed from the seed sequences. The sequence with the highest score from the full group is used for OLG design. The remaining sequences in the full group are used to construct threshold scores used to evaluate the designed OLGs.

### Determining the average success rate from random overlap positions

In order to estimate the expected success rate of an individual overlap attempt, the domains are overlapped at random positions such that one domain is fully embedded into the other. Just as in [30] the sequence with the lower quality of the two constructed OLGs is used as a conservative representative of the pair. After determining the success for each position, the percentage of successful positions for each OLG pair, the average success rate in each reading frame, and the overall success rate averaged across reading frames are calculated. The number of possible positions for each OLG pair is equal to their difference in length plus one, so using more than one overlap position in each pair is only possible for genes with different lengths. Increasing the number of positions for each gene does not change the expected success rate but reduces its variation between different sets (see supplementary Fig. S3). Comparing the variation caused by choosing random positions and the variation caused by choosing random Pfam families, the former turns out to be negligible and consequently only a single randomly chosen position for each OLG pair is used for subsequent analyses. The distribution of the percentage of successful positions in each OLG pair is calculated from up to 50 different positions (see Fig. 4). 50.3% of all OLG pairs form biologically relevant sequences at all positions in every reading frame while only 2.5% cannot form a biological relevant sequence at any position (see Fig. 4). 1.9% of the pairs even form typical proteins, as determined by the 50th percentile threshold, at every position in any reading frame (see right panel in Fig. 3). This result is strongly dependent on the threshold percentile chosen, but due to the wide range of possible results it can still be concluded that the chance of success of a constructed OLG pair depends strongly on the particular genes used, as might be expected.

**Figure 4:**
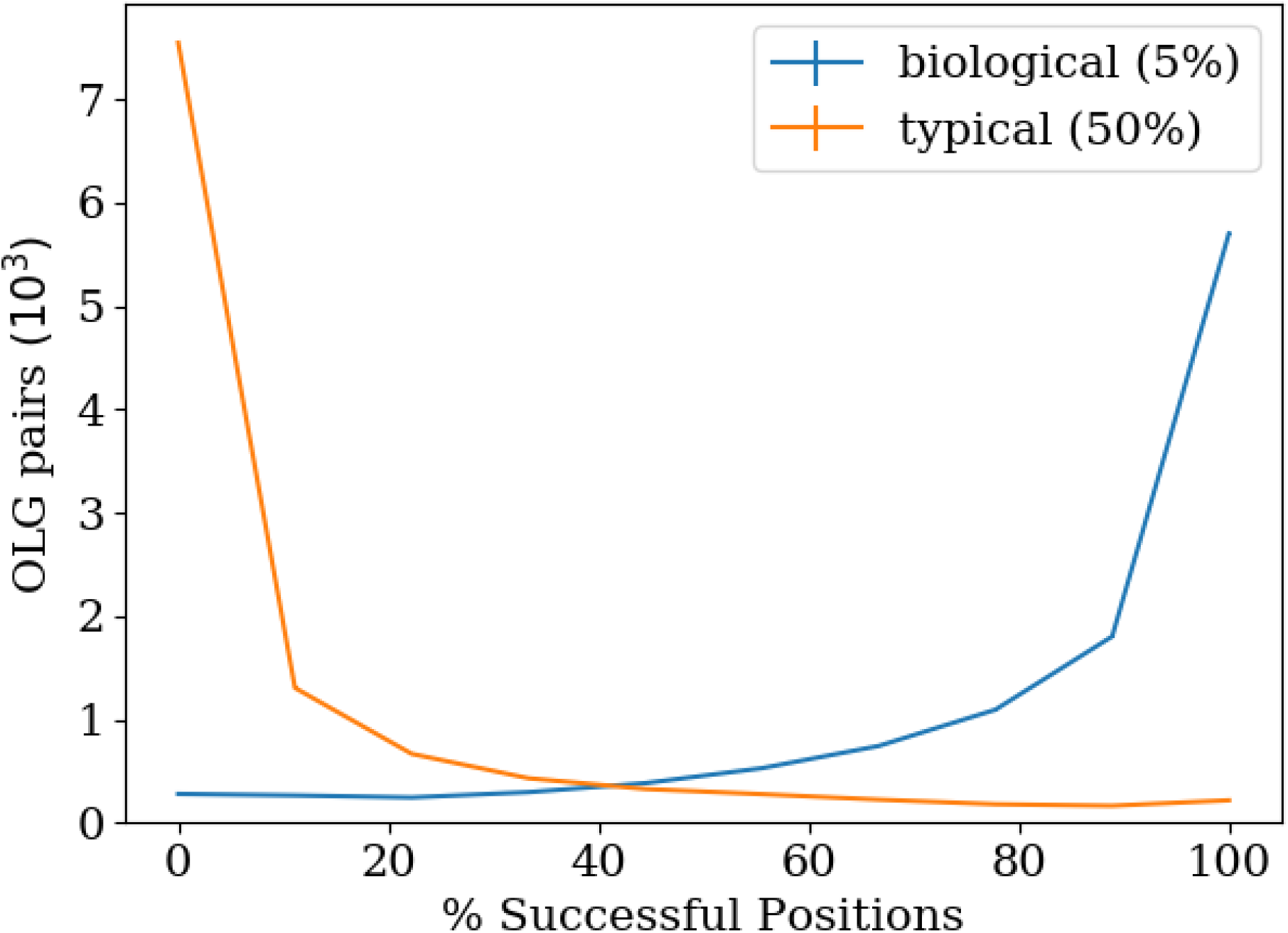
Frequency of successful positions in OLGs. 150 randomly chosen domains with a minimum length of 70 amino acids are used as a basis, resulting in 11325 OLG pairs. In each OLG pair 30 sets of up to random 50 positions were tested against the Pfam group HMMs using the ‘biologically relevant’ threshold (5th percentile) and the ‘typical sequence’ threshold (50th percentile) for a successful overlap. While 50.3% of the pairs can be overlapped at any position and 2.5% in no position using the biological threshold only 1.9% can be overlapped at any position and 66.7% in no position using the threshold of typical sequences. The sequence threshold strongly influences the result.

### Length dependence of the HMM evaluation

In order to determine whether the relative evaluation of OLGs really removed the length dependency, the average quality ‘Q’ of an OLG pair is determined and compared for OLG pairs with different lengths. Q is defined as the ratio of the scores of the constructed sequence (S) over the original sequence (S_max) times 100. The quality is therefore the percentage score loss due to the overlap. Supplementary Fig. S4 shows the mean quality for datasets with different sequence lengths. Starting from around 50 amino acids, Q is indeed mostly independent of sequence length. The low Q values of smaller sequences are because these sequences are less frequently matched to their respective HMM-profile, which results in a score of zero. The reason is probably that the shorter sequences fall below internal detection thresholds of HMMER3 more easily. Changing a single amino acid in a short gene changes its quality to a greater extent than in a long gene, resulting in larger fluctuations, which can lower the sequence below detection thresholds. Lowering internal thresholds of HMMER3 did not lead to more sequences being recognized by their respective profile.

In further analysis the minimum sequence length of 70 amino acids is used so that the percentage of OLG pairs in which at least one sequence is not recognised is below 5% (see supplementary Fig. S4). When taking both sequences of each pair and not only the one with the lower quality, the quality distribution converges to a broad peak at around 76% with increasing sequence length (see supplementary Fig. S5). Since the quality also depends on the flexibility of the HMM profiles used to score the sequences the peak is not expected to get any narrower with increasing sequence length and thus to reduce variations in sequence similarities between the constructed and the original sequences.

### Optimisation of strength of positional weighting

The algorithm to construct OLG sequences from [30] uses an exchange matrix (Blosum62 [41]) to find the closest overlapping sequences to the original ones. It determines the codon with the highest sum of the scores for the exchanges in both sequences at each position. Sequence weights can prioritise the score of either one or the other sequence at different positions in order to increase the chance of obtaining functional sequences. In [30], the weight w_i at position i of the sequence is w_i=e^(-S_i), where S_i is the entropy calculated at position i in the alignment. The weights could be defined differently such that their influence on OLG construction is stronger or weaker. In order to optimize the weight strength a factor k is added to the entropy in their calculation such that w_i=e^(-kS_i). Varying k>0, the optimal weight strength for constructing OLGs can be determined, while k=0 means no weights are being used. In the HMM evaluation the influence of k is very weak. A value of k=0.5 is used in order to maximise the quality, Q (see supplementary Fig. S6). Picking very high k values Q goes to zero. In this case at each position the sequence with the higher conservation maintains its amino acid. This indicates that it is crucial that at each position both sequences are changed in order to create functional OLGs.

In the BLAST evaluation k=0 is optimal (see supplementary Fig. S7), such that no better value can be found for k>0. BLAST does not take special account of conserved regions of a sequence, so weights can improve one sequence but at the same time will reduce the score of the other. Since the lowest scoring of the two sequences is taken to represent the OLG pair, introducing weights has a high chance of reducing the success rate in an evaluation using BLAST, despite increasing biological relevance. This makes an evaluation using HMM or any other method that takes into account sequence conservation significantly preferable for judging constructed OLG pairs.

## Results and Discussion

### Reading frame differences

The five alternative reading frames differ strongly in the combinatorial constraints imposed by the reference gene (mother gene) via the standard genetic code [24], e.g. the sequence N|GCN|, with N being any nucleotide, always translates to alanine in the +1 and the −2 frame. It is interesting whether this difference in constraint transfers to the success rate for designing OLGs. For OLGs resembling typical proteins of their respective families, the success rates for OLG construction varies from 14.9% in the ‘-3’ frame to 3.0% in the ‘-2’ frame with an average value of 9.6% across all reading frames (see Fig. 1). Calculating the e-value just as in [30] as a reference, the constructed OLGs have a median e-value of 10^-(28) to 10^(−37), decreasing with increasing threshold percentile. The result is strongly threshold dependent as 94.5% of the constructed sequences score at least as highly as the worst sequence in the full group, while only 0.02% score better than 95% of the full group. Considering combinatorial restrictions of different reading frames [24] the ranking of frames by success rate are exactly as expected, insofar as the success rate of each reading frame is inversely proportional to the extent of combinatorial restrictions found in [24] (see Fig. 5): the ‘-2’ frame is the least successful reading frame and has the highest restrictions, followed by the ‘-1’ frame, which is the second most restricted frame. Next are reading frames ‘+2’ and ‘+3’, which have exactly the same restrictions and surprisingly almost the same success rates, not only in their average value but also in every single dataset (data not shown), despite expected stochastic fluctuations due to some genes simply fitting better to each other. Last is the ‘-3’ frame, which has no combinatorial restrictions and the highest success rate. Plotting the different success rates in the different reading frames as a function of the number of combinatorial constraints found in [24], results in a linear relation for the lowest possible threshold, namely that all sequences which are at least as good as the worst in the comparison group are judged successful. As the threshold is increased the linear relation is gradually lost (see supplementary Fig. S8). For higher thresholds most of the sequences are below the threshold and very little data is left, which might lead to the observed behaviour. In summary, the structure of the standard genetic code appears to strongly influence the construction of OLGs. Whether the observed relationship between predicted constraints in different frames and the difficulty of constructing OLGs is borne out by the proportion of natural OLGs found across frames deserves attention across diverse taxa.

**Figure 5:**
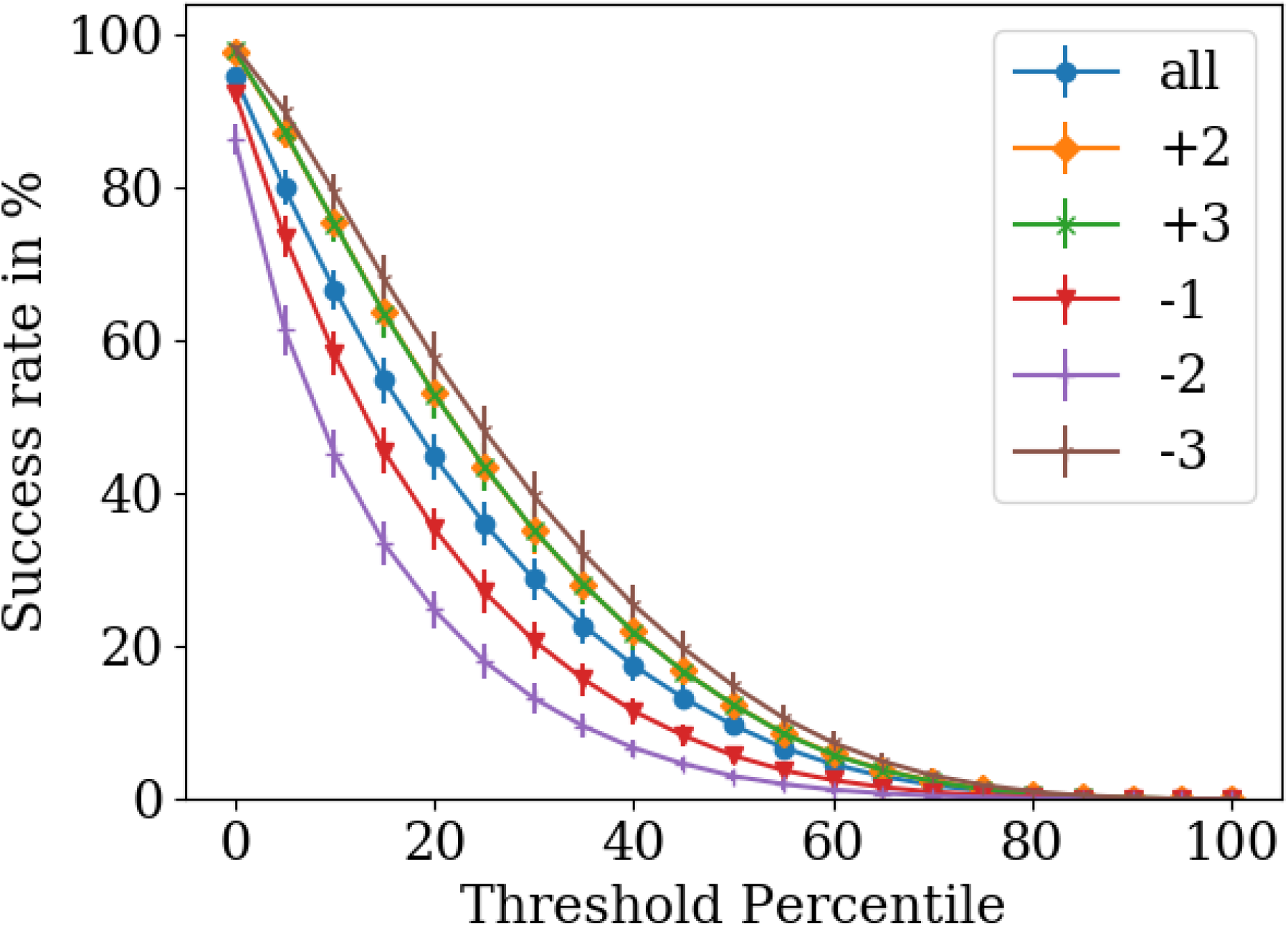
Success rates for OLG design in different reading frames as a function of threshold percentile. Each value is an average from 20 different datasets of 150 sequences with at least 70 amino acids and the error bars are equal to the standard deviation. The threshold chosen within the Pfam group has a very strong influence on success rates. The ordering of the reading frames by success rates, namely ‘-3’, ‘+2’/’+3’, ‘-1’ and ‘-2’, matches the ordering by combinatorial restrictions in the standard genetic code, beginning with the least restricted frame [24].

### The impact of OLG design on amino acid sequences

Determining the impact of OLG construction on an amino acid sequence identity is another indicator of its functionality. It has been argued that a 34% amino acid identity between naturally occurring sequences ensures that both sequences have the same structure [54]. Comparing the altered part due to OLG construction with the original sequence, in 96.5% of cases both OLG sequences share at least 34% of amino acids with their original sequence. In some OLG pairs both sequences have an amino acid identity of up to 60% compared to their original sequence. In the biologically more relevant property of amino acid ‘similarity’, the worst-scoring of the two OLGs can be even up to 80% similar to its respective original sequence (c.f. left panel of Fig. 6). Determining the average amino acid identity and similarity between the two OLG sequences, the average OLG design impact can be determined. The average amino acid identity is 60% in most cases (right panel of Fig. 6) showing that in almost all OLG pairs one sequence is above and one is below 60% amino acid identity. The average amino acid similarity is 75% in most cases (right panel of Fig. 6) which again shows that in almost all cases one of the two OLG sequences is above and one below 75% identity. The double peak structure of both panels in Fig. 6 can be explained by differences for OLG pairs in different relative reading frames, which are pooled here (c.f. Supplementary Fig. S14). It follows that in an average OLG design, in 20% of all overlap positions the amino acids of both sequences can be maintained, in 30% one sequence maintains its amino acid while the amino acid in the other sequence is changed to a similar one and in 50% one sequence maintains its amino acid and the other sequence cannot maintain a similar amino acid. How well the two sequences can be maintained after the overlap is determined by the standard genetic code and the two specific sequences, the overlap position, their amino acid composition and the amino acid order. While the standard genetic code is a constant factor across all overlaps, all other factors are specific in each case and create the variability in the results.

**Figure 6:**
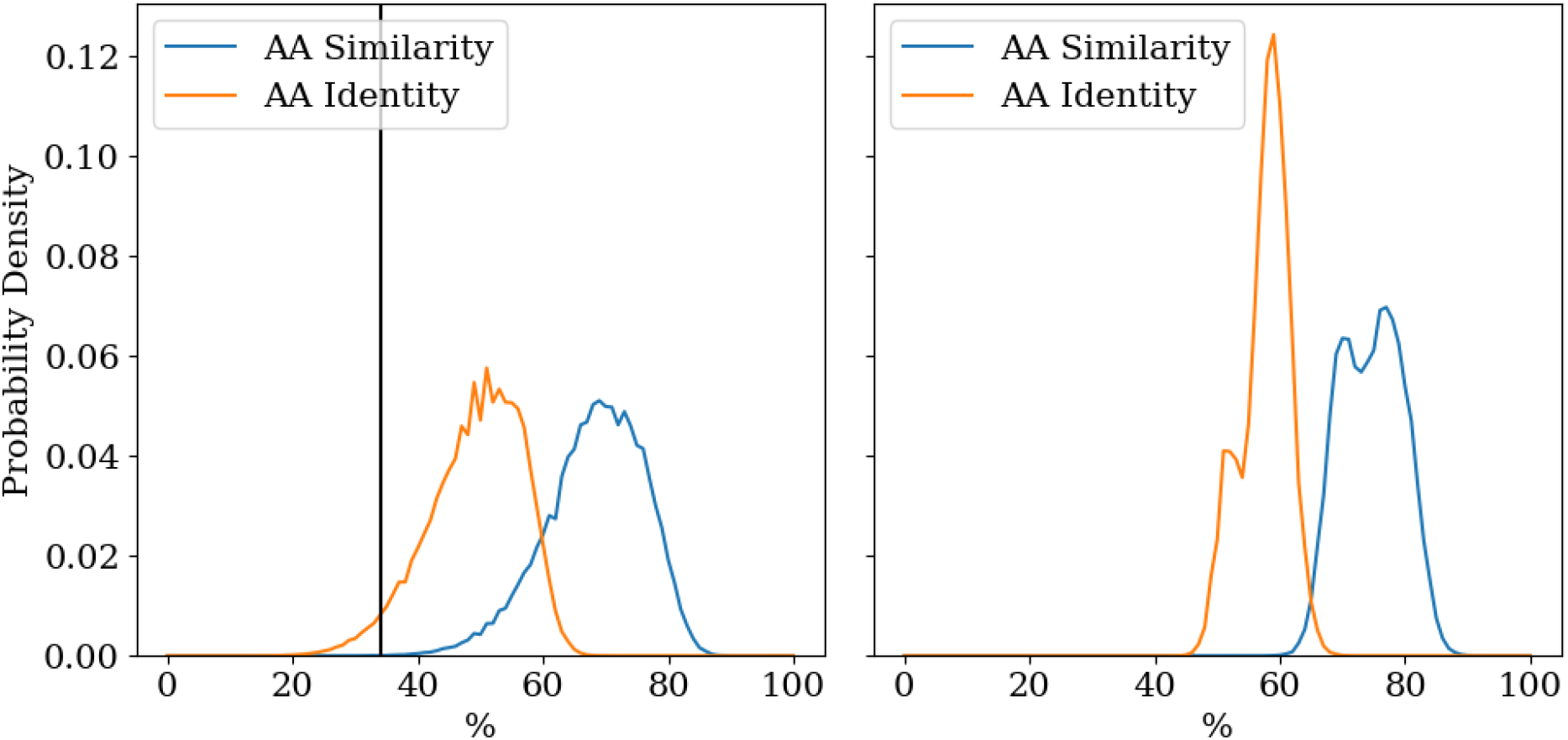
Probability density for different amino acid identities and similarities in constructed OLG pairs. The data is calculated from 505,000 OLG pairs. *Left:* The sequence with the lower identity is representative of the pair. The black line indicates the 34% amino acid identity threshold. *Right:* The mean similarity of both OLG sequences represents the pair.

The impact of OLG design on secondary structure is the last factor studied here. Comparing the secondary structure of the OLG sequence with its original sequence, a secondary structure similarity is determined. Secondary structure is predicted using Porter 5 [42] with the “--fast” flag. It can distinguish between the eight different secondary structure motifs of the dictionary of protein secondary structure (DSSP) [47,48,49], which are 3_10-, alpha-, and phi-helices, hydrogen bonded turns, beta sheets, beta bridges, bends and coils. Determining the same secondary structure similarity for all sequences in the seed group of the Pfam database yields a control group. This way the typical deviations between domains with the same function can be determined. Comparing probability densities for different secondary structure identities in both groups it can be seen that the constructed OLG sequences barely deviate from the seed sequences (c.f. Fig. 7). In conclusion, in regards to secondary structure the change inflicted on a sequence to create OLGs is no more than the differences within naturally occurring protein domain families.

**Figure 7:**
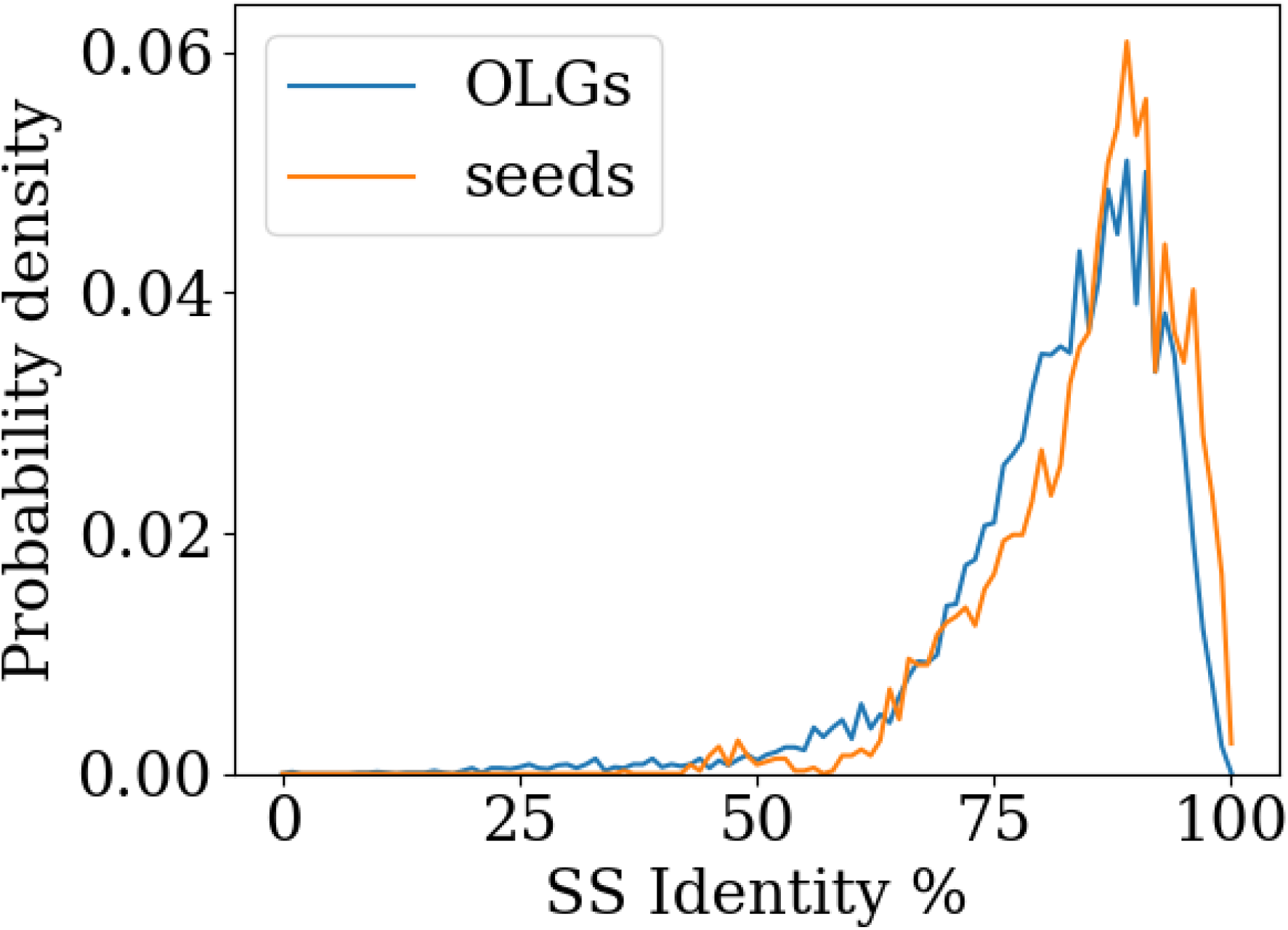
Probability densities for different secondary structure identities for OLGs and seed sequences calculated from a dataset of 50 sequences consisting of at least 70 amino acids. OLGs are as similar to their original sequences in secondary structure as observed for comparisons of seed sequences of naturally occurring protein domains to the sequence best representing the respective domain family.

It is noteworthy that only amino acid identity and similarity have a strong correlation (r=0.82) so combined with the other parameters, namely the relative HMM score and the secondary structure identity, there is a set of three more or less independent properties for evaluating constructed OLGs, and probably for protein homologs in general. The relative HMM score is the HMM score of the OLG sequence divided by the HMM score of a sequence at any threshold percentile as discussed above. Between each pair of parameters the Pearson’s correlation is below 0.2, with the exception of the correlation between secondary structure identity and HMM score being r=0.37 or r=0.39 for thresholds of 95% or 100% respectively.

### Influence of the genetic code and code optimality

By comparing OLG sequences constructed with the standard genetic code (SGC) to sequences constructed with artificial codes the level of optimality of the SGC can be inferred. Since such an approach depends strongly on the codeset used [43], four different versions with increasing restrictions will be tested. There are two factors defining a genetic code, namely its amino acid composition and the arrangement of amino acids on the 64 codons. The first code set is the random code set and does not constrict any of the two factors. Each code can have any of the 20 amino acids used in the SGC at any codon. The second set only restricts the composition of its codes and is called the degeneracy code set. All codes in this set contain the same amount of codons for each amino acid as in the SGC and thus conserving its amino acid composition. The third set is the blocks code set whose codes have a very similar structure to the SGC and while it also restricts the composition of the codes to some degree it mostly determines their arrangement. This code set is created by assigning all codons of the SGC that code for the same amino acid into blocks and shuffling the amino acids assigned to each block and thus conserves the degeneracy structure of the SGC on the third nucleotide. Lastly a code set that maintains the mutational robustness of the SGC as calculated in [43] is tested. In short, the mutational robustness is the average change of amino acids due to point mutations and has been shown to be extremely optimal in the SGC relative to similar codes [44]. This set contains block codes like in the second set but only the codes whose mutational robustness is at least as high as the SGC are kept. Since these codes are fundamentally block codes they are partly restricted in their amino acid composition, but the arrangement of amino acids in these codes is even more restricted as point mutations from any codons should result in similar amino acids. This code set reflects the fact that different properties of the SGC have a different impact on the fitness or biological optimality of the SGC with the mutational robustness most likely being one of the most important features. Here this code set is called the mutational robustness blocks set (MR-blocks set) and it tests the optimality of constructing OLGs as an additional property of the SGC after taking into account the mutational robustness.

Comparing the degeneracy, the block and the MR-blocks code set to the random set, the influence of code composition and arrangement can be determined (see left panel of Fig. 8). The degeneracy code set reflecting the composition of the SGC has the codes with the highest average success rates indicating that the composition of the SGC is a major factor for this property, but the SGC itself has a very low success rate in comparison, indicating that the amino acid arrangement is an even stronger - in this case negative - factor as the SGC is worse than both the random codes and the degeneracy codes. The block structure of the SGC has a strong negative impact on successful OLG design and the SGC is a typical member of this set. Enforcing even more structure on the artificial codes in order to maintain the mutation robustness of the SGC further reduces the ability of the SGC to create successful OLGs.

**Figure 8:**
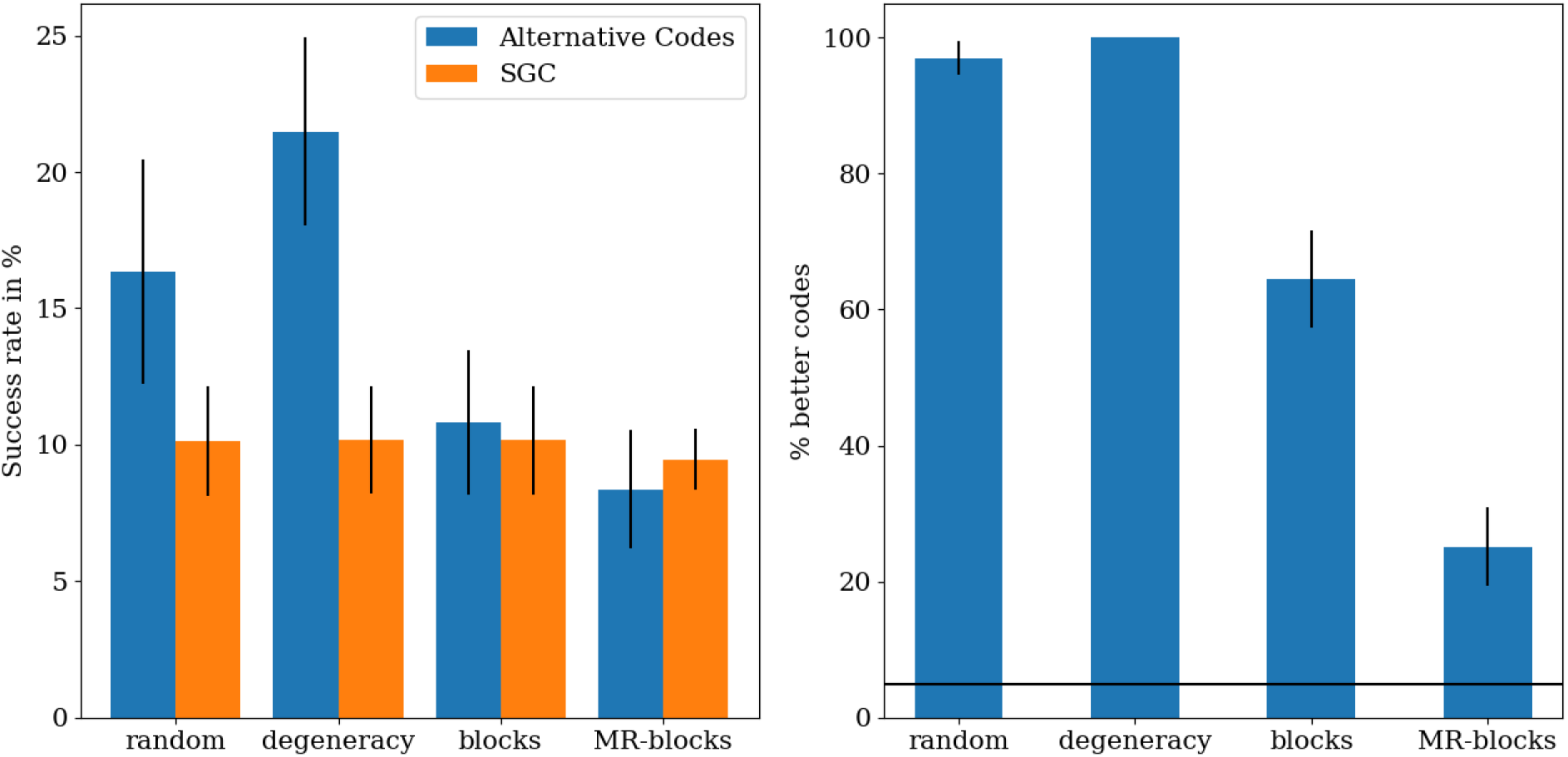
OLG design success rates for different alternative code sets. The average is calculated from 20 sets of 100 alternative codes, except for the MR-blocks set with 10 sets of 500 codes. The error bars indicate the standard deviation. *Left:* The average success rates compared to the SGC. While the composition of the SGC is a positive factor, the arrangement of the SGC is a negative factor. *Right:* The optimality of different code sets. The black line indicates the 5% threshold. The more restricted the code set the more optimal the SGC appears indicating that the ability to successfully create OLGs has only been optimized while maintaining other properties.

Studying the optimalities of each of the four code sets for flexibility in OLG design, it is apparent that the more restricted the code set is, the more optimal the SGC is relative to the set (see right panel of Fig. 8). Especially in the MR-blocks code set only a few codes are better than the SGC, however no codeset or reading frame has fewer than 5% of codes doing better (see Fig. S19-S12), which has been a recommended threshold for inferring optimality [45]. This is an expected result even if the code has been optimised for OLGs as the success rate for constructing OLGs reflects merely the ‘flexibility’ of a code system, but OLG sequences also need to be conserved, which is an almost directly opposing property which also has not been found to be strongly optimal by itself [43]; it might indeed be expected that overall optimality involves a trade-off between the two.

If the SGC has been optimized in this way this could indicate a turning point at which a further increase in mutational robustness results in a smaller fitness increase compared to an increase in the flexibility to create OLGs - how to measure fitness for a genetic code is however not clear. While the code composition of the SGC is beneficial for both the ability to create successful OLGs and the mutational robustness, the code arrangement of the SGC is only beneficial for mutational error robustness and the SGC (see Fig. 2 of [43]), indicating that the mutational robustness is the more important property. Only in the set of codes with the same mutational robustness the optimality for OLG design becomes stronger, supporting the turning point hypothesis.

### Taxonomic differences

Besides the four basic taxonomic groups (three domains of cellular life: archaea, bacteria, eukaryotes, plus viruses) also old genes can be studied by picking only families which have at least one sequence in all four taxonomic groups since it is expect that these families have already been present in LUCA or another ancient ancestor (although this high level categorisation is not perfect due to widespread horizontal gene transfer). Surprisingly, bacterial and eukaryotic genes are generally significantly better suited to OLG construction than virus and archaeal genes with only minimal dependence on the threshold percentile, c.f. Fig. S13. The largest dependence on the threshold percentile is found for the “Found in All” genes, which only a total of 50 sequences can be found in the Pfam database, so higher stochastic fluctuations are to be expected. Using the ‘biologically relevant’ threshold, the biggest difference is between eukaryotic and archaea genes which have a 20% difference in their success rate (see left panel of Fig. 9). For OLGs which are typical proteins of their respective family, eukaryotic genes are almost twice as likely to be successful as virus genes (see right panel of Fig. 9).

Eukaryotes and “Found in All” genes are typically the easiest to overlap, which is somewhat unexpected as eukaryotic genes would perhaps be expected to have the youngest protein families, and so to appear less ‘flexible’ due to having sampled less of the functional space through mutations. More understandable however is that due to being closer to mutational saturation (if more ancient on average) and therefore having explored a larger proportion of functional sequence space, “Found in All” genes might appear more ‘flexible’, resulting in lower weights and thresholds.

**Figure 9:**
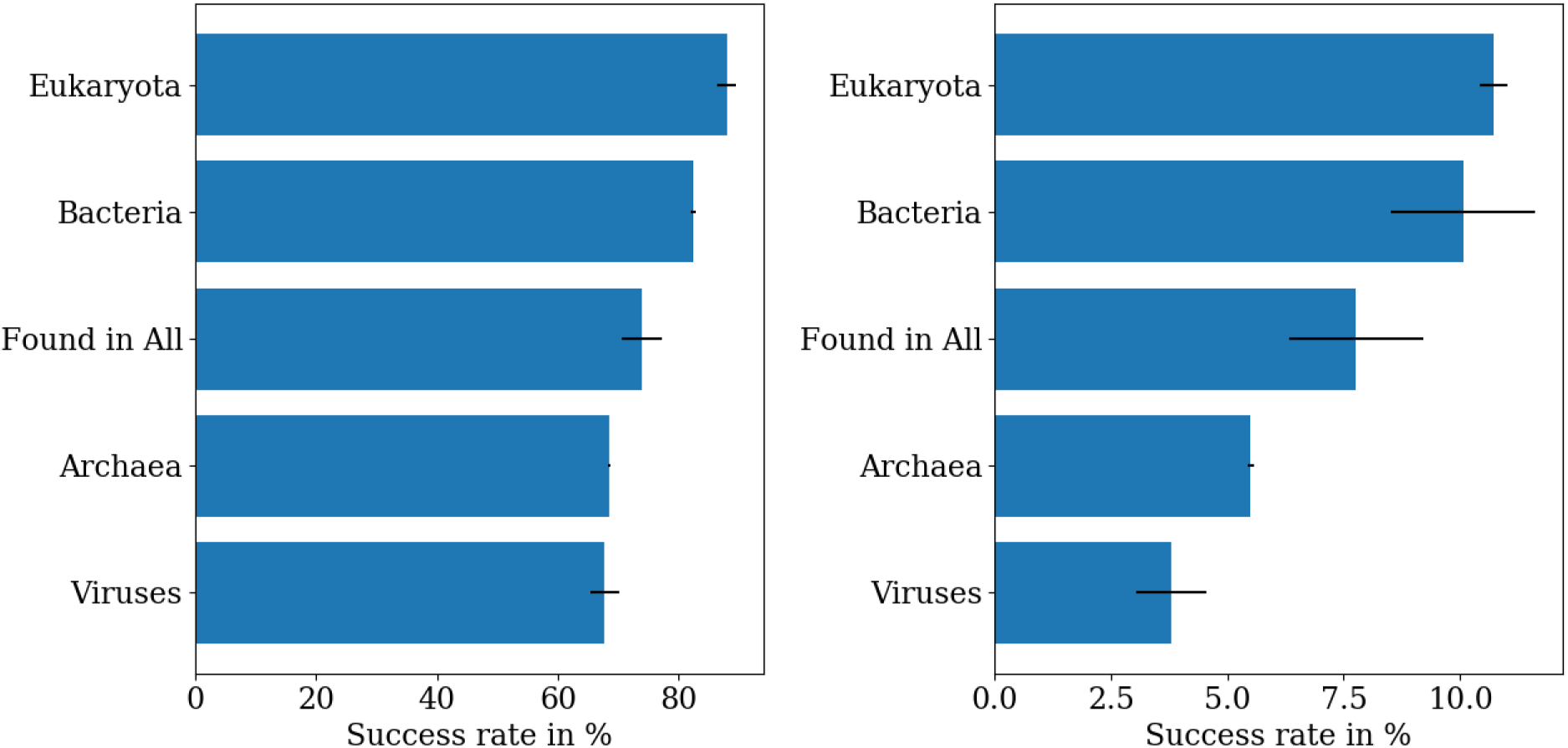
Average success rates in different taxonomic groups calculated from 20 datasets with 150 domains consisting of at least 70 amino acids each; the black lines indicate the standard deviation. The ordering of ‘biologically relevant’ genes (threshold percentile 5%, *left*) is mostly identical to ‘typical’ genes (threshold percentile 50%, *right*). Sequences from Bacteria and Eukaryota sequences outperform Viruses and Archaea.

### Evolutionary distance of OLGs to biological sequences

In order to estimate the difficulty of naturally forming OLG sequences, the minimum number of nucleotide changes needed in order to reach the OLG sequence from any of the two original sequences is determined (see Fig. 10). By only taking OLGs in which both sequences are above a certain HMM threshold, extreme outliers are gradually removed with increasing threshold but the rest of the distribution stays the same. This indicates that this property is independent of the threshold value, just as for the amino acid identity and similarity, as fewer and fewer designed OLGs pass a higher threshold which makes extreme outliers less likely to occur. On average a designed OLG sequence has a 25% difference in nucleotides to its original, with half of constructed sequences in the range of 20-30% change. Most interesting are outliers on the lower end of the distribution as they indicate whether OLGs exist that are potentially reachable by naturally occurring mutations. The lowest nucleotide difference observed is 1.8%, which was for an OLG pair that scores better than 25% of the domains in the comparison group. 0.6% of OLGs required less than 10% nucleotide change, i.e. 5843 sequences of the 955846 sequences created in this dataset that scored at least as highly as the worst sequence in the comparison group. This suggests intuitively that creating overlaps of the sort constructed here could be possible naturally through accumulation of random mutations. The population genetics of such a hypothetical process is a potential topic for further study, as is an experimental evaluation of functionality.

**Figure 10:**
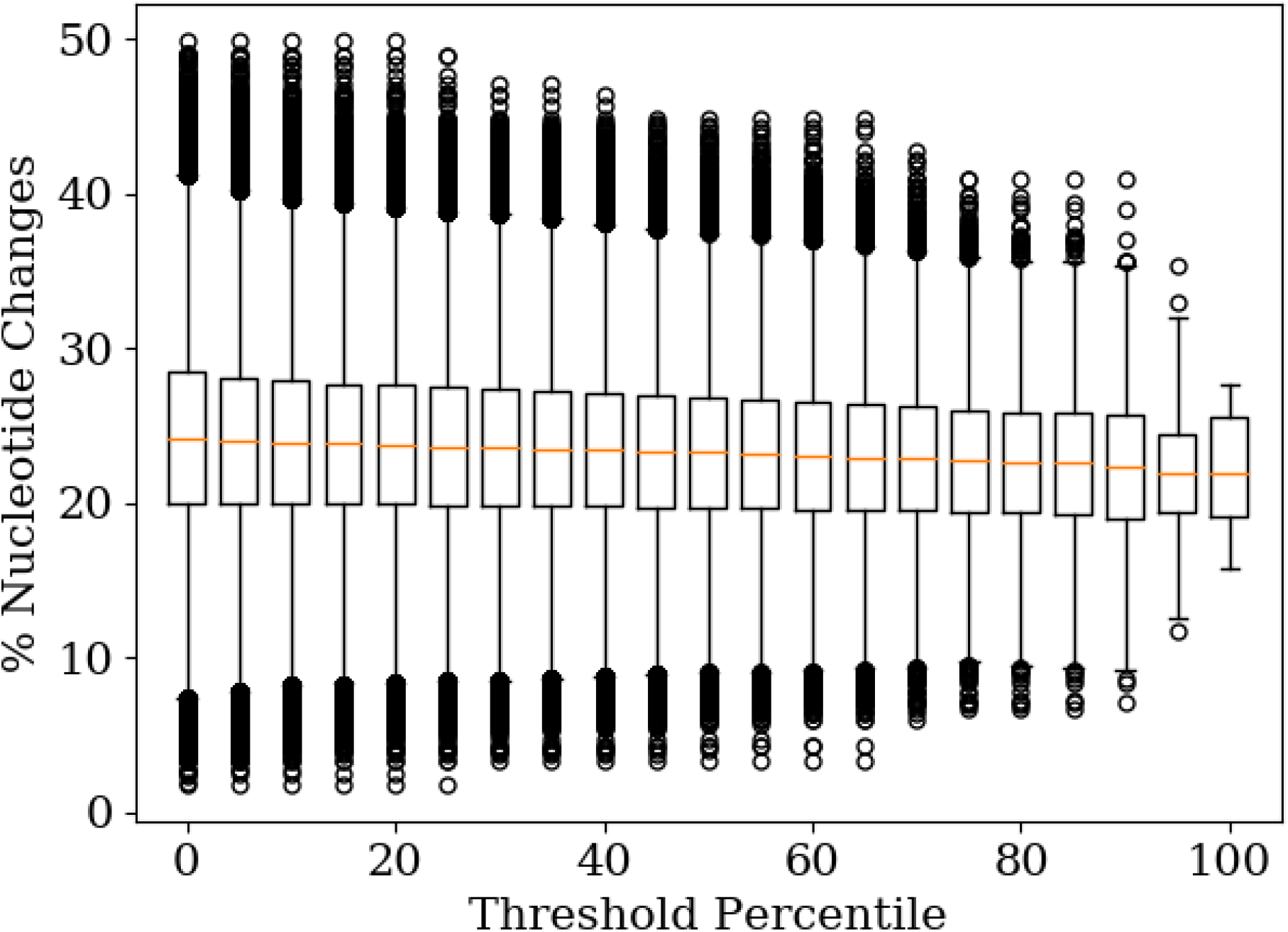
Percentage nucleotide change of OLGs as a function of HMM threshold %. The minimal nucleotide distance of each of 1010000 OLG sequences (two per pair), with a minimal length of 210 nucleotides, to their respective original sequence is determined.

## Discussion

There are many aspects of the synthetic construction of OLG pairs which can be studied. Here factors such as sequence length and the influence of sequence conservation are taken into account. The analysis shows that an evaluation with BLAST and a fixed e-value cutoff cannot accurately assess the potential functionality of the designed OLGs. While the combination of sequence length and an e-value cutoff completely determines the success rate of the constructed OLGs, adding in positional weights can only negatively influence the sequences constructed with this method. Both problems can be solved however by instead using HMM profiles to determine sequence similarity and then using these to define a threshold for successful OLGs derived from sequences in the same protein family. The HMM profiles and the thresholds are though both derived from the Pfam database [38], which makes these results strongly dependent on the database quality. For example, if in one taxonomic group sequences are very similar due to being mostly from the same species or genus, thresholds would appear to be higher and it would be harder for designed OLGs to pass these thresholds. Further optimization of the construction algorithm can be achieved by determining the optimal weight strength (influence of sequence conservation), which is k=0.5.

94.5% of the constructed OLG sequences score at least as highly as the worst-scoring biological sequences in Pfam groups, while 9.6% of the sequences cannot be distinguished from naturally occurring domains in their respective protein family. This indicates that the typical variation inside protein families is of the same order of magnitude as the change needed in order to construct artificial OLGs by arbitrary pairing of protein domains. This result also holds true for other bioinformatic factors like amino acid identity and secondary structure, since the constructed OLGs are typically very similar to naturally occurring domains in these properties. Studying artificial OLG design success from the perspective of an even more constricting biological parameter like tertiary structure would be an important next step; but besides the amino acid sequence, also codon usage can impact protein structure [52], along with environmental factors such as the presence of chaperone proteins, which together make it a much harder problem. Ultimately, proof of the functionality of artificial sequences cannot yet be realised bioinformatically, and experimental verification is essential. To this end all known independent protein properties available from the sequence should be tested in order to create a gold standard for possibly functional sequences. From this study it is clear that sequence similarity (or identity), HMM-scores and secondary structure should be part of the judged properties. Determining relative HMM scores for high thresholds could be used to prefilter sequences for secondary structure prediction as it is the computationally most intensive part of this analysis.

Considering that domain-domain overlaps are expected to be much harder than overlapping a domain with a less conserved region in another gene, it appears that de novo origin of genes from overlapping ORFs may be much less difficult than widely assumed. Some constructed OLG sequences varied only by 1.8% from their original sequence, and there might be other natural sequences from the same domain that are even closer to the OLG sequence. This result could be a starting point for estimating the difficulty of evolving OLGs from different starting sequences in natural systems, which is still relatively unexplored despite some early work [46].

The structure of the standard genetic code explains differences between reading frames and is a strong factor in the overall success rate of OLG construction. OLGs can maintain an average 60% amino acid identity and an average 75% amino acid similarity, which is mostly due to the genetic code. The structure of the standard genetic code is defined by its composition, namely how many codons code for each amino acid, and its arrangement, namely which codons code for each amino acid. It is known that the composition alone can not explain the strong optimality of the standard genetic code for mutational robustness as it stands out from between codes with the same composition as the standard genetic code [43,55]. Considering that the arrangement of the standard genetic code creates such high mutational robustness values [44] it is remarkable that designing OLGs also works so well.

Another factor which deserves further exploration is the age of a protein family, i.e. the time since gene birth. This may correlate with apparent ‘sequence flexibility’, which is the strongest influence on the result via the threshold values, due to increasing mutational saturation in older protein families. Being able to distinguish genuine sequence flexibility from mutational saturation, even in broad terms, could be very useful here.

The analysis presented here depends primarily on the reliability of HMM profiles of Pfam groups as a guide to biological functionality in constructed sequences. Reliability for classifying biological protein sequences into ortholog families, the main use of these HMMs, may not correlate well with reliability in scoring artificially constructed sequences for functionality. In other words it may well be that these profiles fail to capture important requirements for protein tertiary structure and/or functionality. Future research ought test the best candidates experimentally, and if the best candidates from the methods developed here are not successful, additional factors could be considered in comparing constructed sequences and their natural precursors. For instance, many protein characteristics can be assessed using servers or packages incorporating multiple bioinformatic tools such as PredictProtein, for various secondary structural elements [57], and many sequence properties, such as hydrophobicity profiles, can be computed using the VOLPES server [58], which has been applied to the related case of frame-shifted sequences compared to their mother genes [16]. Other properties required for functional protein sequences can be inferred from the evolutionary information contained in sequence alignments of protein families. For instance, it has been calculated based on a study of residue-residue co-evolution in ten well-characterized protein families that the proportion of all sequences which fold to the family’s structure ranges from approx 10^-24^ to 10^-126^ [59]. These principles have recently been successfully used in the design of functional proteins [60], and could conceivably also be applied to OLG construction.

Factors facilitating the existence of OLGs may possibly help in predicting OLGs in sequenced genomes and should be explored further. For instance, a careful study of relatively ‘flexible’ sequence regions in taxonomically widespread genes may help find more overlapping genes. Most interestingly, bacterial and eukaryotic genes can be overlapped more easily than virus genes, contrary to the findings in [30]. These earlier results can be explained entirely with dataset-database biases, so this algorithm gives no support for the common assumption of a higher intrinsic OLG formation capacity of viruses compared with bacteria or eukaryotes. Two of the main differences between the taxonomic groups are the expected mutation rates and the average length of a protein. While genomes with higher mutation rates explore sequence space faster and therefore their proteins should appear to be more flexible, virus domains do not appear to be very flexible, despite having the highest mutation rate. The length of the sequences on the other hand has been removed as a factor in this analysis. An artificial factor not considered could be database biases or an exchange matrix (BLOSUM62) biased towards certain kinds of proteins. The latter could be tested by using different matrices created from sequences from different taxonomic groups. It would be important to use the new matrix not only in the construction of the OLGs but also in the evaluation by the HMMs. So far it is not clear why protein families from different taxonomic groups are so different in their calculated ability to create OLGs. A better theoretical understanding of overlapping genes will be extremely useful in microbial genome annotation methods, the study of evolution, and in synthetic biology, and therefore deserves renewed attention.

